# The MLR COMPASS-like complex is required to regulate triglyceride depletion and stress response in the Drosophila fat body

**DOI:** 10.1101/2021.04.01.438109

**Authors:** David J Ford, Claudia B Zraly, Jeffrey Ng, Andrew K Dingwall

**Affiliations:** Department of Cancer Biology, Stritch School of Medicine, Loyola University Chicago, Maywood, IL, 60153, USA; Department of Pathology & Laboratory Medicine, Stritch School of Medicine, Loyola University Chicago, Maywood, IL, 60153, USA

**Keywords:** Enhancers, Drosophila, Metabolism, Triglycerides, Stress Response

## Abstract

MLR COMPASS-like complexes are highly conserved histone modifiers and enhancer regulators recruited by multiple transcription factors during differentiation and development, including lineage-determining factors, nuclear receptors, and developmental signaling pathway effectors. While the essential functions of MLR complexes during differentiation-associated transcriptional reprogramming has been well-explored, roles that these complexes play during reprogramming events in terminally differentiated cells remain understudied. We determined that the Drosophila MLR complex is required in fat body adipocytes for proper regulation of the triglyceride depletion rate during non-feeding periods, including metamorphosis and adult starvation. Transcriptome analysis revealed that the complex plays a critical role in controlling stress-related reprogramming in these cells, suggesting that the metabolic phenotypes are indirect effects of dysregulated stress transcription. Furthermore, our evidence suggests that the complex interacts with Foxo and Relish (Nf-κb) pathways and is required for proper expression of their targets. This investigation further elucidates the necessary functions of MLR complexes in regulating transcriptional reprogramming in terminally differentiated cells as well as suggests novel binding partners. Apparent roles for these complexes in the proper regulation of stress-induced transcription implies new mechanisms involved in cancer and other human diseases associated with MLR subunit mutation.

## Introduction

COMPASS-like complexes (<u>Com</u>plex of <u>P</u>roteins <u>As</u>sociated with <u>S</u>et1) are ancient and highly-conserved histone modifiers responsible for methylation of the lysine 4 residue of histone 3 (H3K4), an epigenetic mark associated with open and active chromatin ^1^. Multicellular organisms contain multiple paralogous COMPASS-like complexes specialized for specific genomic targets and methylation quantity; while several subunits are common among complexes, each individual complex is characterized by a unique methyltransferase subunit. Those containing methyltransferases KMT2C (MLL3) or KMT2D (MLL2/4) in humans and Trr in Drosophila belong to a branch known as MLR (<u>ML</u>L/Tr<u>r</u>) complexes ^2–4^. During transcriptional reprogramming events, MLR complexes are recruited to distal regulatory enhancer regions by a variety of transcription factors or other binding partners. Enhancers promote full activation and efficient transcription of their target genes, which is accomplished in part by physical contact with target promoters ^5,6^. Once bound, MLR complexes catalyze the deposition of H3K4me1, contribute to the removal of H3K27me3, and are required for recruitment of the histone acetyltransferase p300/CBP, all of which are necessary steps for full enhancer activation ^7–10^. While understandably recognized as co-activators of enhancers and their downstream transcriptional activity, recent investigations by our group have characterized multiple examples of repressive activity by an MLR complex, suggesting a role in enhancer suppression for the prevention of premature activation ^3,4^. Proper enhancer regulation, and therefore MLR complex activity, is required for the elaborate patterns of spatiotemporal transcriptional reprogramming that occurs during multicellular development; alteration of MLR complex activity results in defects in tissue patterning, lineage determination, and cellular differentiation^10–14^. For example, murine Kmt2c and Kmt2d colocalize with adipogenesis factor Pparγ and are required for proper fat tissue development ^14,15^. Transcriptional reprogramming events also occur subsequent to differentiation in terminally differentiated cells, often in the context of response to extra-cellular stimuli. Potential roles for MLR complexes in these types of reprogramming events in an intriguing yet understudied area. It has been previously determined that MLR complex activity is necessary for critical digestive bile acid transporter expression in hepatocytes in response to homeostatic signals ^16,17^. In order to continue the investigation and identify further examples of MLR complex regulation of reprogramming in terminally differentiated cells, we turned to the model organism Drosophila, the fruit fly.

*Drosophila melanogaster* contains a single tissue type homologous to both mammalian adipose and hepatic tissue: the fat body. While the fat body varies in structure and size depending on developmental stage, it maintains its role as a critical regulator of systemic metabolism, developmental signaling, and stress response ^18^. Its functions include synthesis and mobilization of triglyceride (TAG) droplets for energy storage, regulation of homeostatic responses to feeding or non-feedings states, and secretion of antimicrobial peptides as a part of the innate immune response ^19–24^. The antagonistic balance between feeding/growth and stress response states is critical for animal survival, and through its manifold functions the fat body acts as a master regulator of this balance.

During feeding/growth periods, fat body adipocytes uptake metabolites from digestion and convert them into forms of stored energy: glycogen and TAGs. These collected TAGs act as the main source of metabolic energy during non-feeding periods, including metamorphosis and starvation ^20,25^. TAGs are depleted mainly by lipase Bmm, which converts diacylglycerol for shuttling into the hemolymph for systemic usage ^19^. Transcription of Bmm and other TAG-regulating machinery is controlled by mutliple antagonistic signaling pathways sensitive to feeding and stress states ^26^: TAG storage is promoted by insulin-like peptide induction of Akt signaling during feeding ^27^, TAG depletion is promoted by Akh and steroid hormone ecdysone signaling during starvation and developmental transitions ^28–30^. In addition to the direct activation of nuclear receptors and other transcriptional regualtors, these metabolic signals indirectly regulate stress response through master stress factor Foxo, which acts as the fulcrum of the axis balancing healthy homeostasis and stress response ^20,31–33^.

In terminally-differentiated cells, stress response genes are kept is a primed state of paused transcription initiation, similar to the poised chromatin architecture awaiting developmental signals in undifferentiated cells ^34,35^. Transcription of many of these critical protective genes relies on Foxo activity ^36–38^. While large metazoans harbor multiple specialized Foxo paralogs, Drosophila contains a single Foxo factor responsible for stress response regulation in a tissue types ^39,40^. In addition to common stress-response genes, Foxo has adipocyte-specific regualtory targets in the fat body, including Bmm and antimicrobial peptides (AMPs). Foxo is the primary direct activator of Bmm expression and therefore TAG depletion rate during periods of starvation ^41,42^. Precise modulation of the TAG depletion rate to both ensure survival and avoid premature exhaustion of energy stores is accomplished through NF-κB factor Relish, which acts antagonistically to Foxo and supresses Bmm expression during starvation ^43^. Both Foxo and Relish are also responsible for the regulation of AMP expression in adipocytes, which are secreted by the fat body in response to microbial invasion to damage the foreign baterial or fungal cells ^44,45^. While conventionally regulated by Toll or Imd pathway effectors such as Relish during antimicrobial stress, AMP transciption occurs independently of innate immune signaling during periods of developmental change, such as metamorphosis ^46–48^. AMP expression in these latter situations requires Foxo ^48^.

In this study, we investigated functions of the Drosophila MLR COMPASS-like complex in the fat body by knocking down or overexpressing essential subunit Cmi exclusively in adipocytes. We determined that while enhancement or depletion of complex activity has no effect on post-embryonic fat body development and TAG synthesis, complex activity is required to precisely regulating the rate of TAG depletion during metamorphosis and adult starvation. Transcriptome data suggested the MLR complex is required to regulate stress-response transcription, potentially through interaction with Foxo and/or Relish. Our results suggest that the MLR complex is required for transcriptional reprogramming in terminally differentiated cells, specifically poised metabolic/stress-response genes in adipocytes, and that this control is necessary for proper metabolic response during non-feeding periods.

## Materials and Methods

### Drosophila Husbandry and Stocks

All stocks and genetic crosses were reared on standard cornmeal/dextrose medium (13% dextrose, 6% yellow cornmeal, 3% yeast extract, 1% agar, 0.3% methylparaben) at 25°C. Experimental crosses were performed at 28°C unless otherwise specified. UAS-Cmi-IR and UAS-Cmi lines were previously described ^49^. Other fly strains were obtained from the Bloomington Drosophila Stock Center (BDSC), including Lsp2-Gal4 (#6357), UAS-Foxo-IR (#32427), and UAS-Foxo (#80946). These strains are described in Flybase (http://flybase.bio.indiana.edu).

### RNA-sequencing

Wandering third instar larvae were collected and dissected in ice cold PBS. Entire fat body tissue from twenty-five larvae was extracted, ensuring no collection of other tissue types, and placed in ice cold PBS. Each collection was flash frozen with dry ice until preparation of tissue for RNA extraction. DESCRIBE RNA EXTRACTION. RNA-seq and statistical analyses was performed by Novogene Genomics Services.

### Survival Analysis

To track metamorphosis survival, wandering third instar larvae were placed in vials and allowed to undergo metamorphosis at either 25°C, 29°C, or shifted between the two temperatures at a midpoint of metamorphosis. For temperature shift experiments, tubes containing pupae were transferred from the initial to the secondary temperature at approximately 50 hours post-pupariation (∼stage P7 according to Bainbridge and Bownes pupal staging) ^50^. Survival was recorded by tallying number of living newly-eclosed adults per number of larvae placed in each vial. “Pharate Lethal” was defined as a fully formed pharate in the pupal case that is not moving or beginning eclosion (stage P14 according to Bainbridge and Bownes pupal staging) and is by all appearances dead, not responsive to stimuli or to removal from pupal case. The experimental and control animals were staged and placed in parallel in all cases. “Eclosion Lethal” was defined as a fully formed adult in the process of eclosion from the pupal case (stage P15(ii) according to Bainbridge and Bownes pupal staging) and is by appearances dead, not responsive to stimuli or to removal from pupal case.

To track adult starvation survival, newly-eclosed adults (0-5 hours post-eclosion) were transferred to vials containing starvation media (2% agar and 0.25% methylparaben antifungal) to provide moisture but no carbohydrates, amino acids, lipids, or other nutrients. Survival was assayed by tallying clearly living flies (assayed by movement) at regular timepoints until death of all individuals. The experimental and control individuals were starved and assayed in parallel in all cases. Results were plotted as survival probability according to Kaplan-Meier estimation and statistical difference between control and experimental populations was measured using the log-rank test. For aged animal experiments, newly-eclosed adults were separated according to sex and kept on standard media for 4 days to allow for TAG levels to reach adult homeostasis. Animals were then transferred in parallel to starvation media and survival was assayed similarly. For refeeding assay, newly-eclosed adults were transferred to starvation media for 24 hours, at which point they were transferred onto standard medium. For use in TAG quantification, three samples of each genotype were collected immediately after eclosion, after 1 day starvation, after 1 day refeeding, and after 2 days refeeding. Each replicate consisted of five animals homogenized.

To track oxidative stress survival, oxidative stress media was prepared by adding to the standard food recipe hydrogen peroxide (H2O2) to final concentration of 5%. Newly-eclosed adults were separated by sex and aged for three days before being placed on starvation media for 4 hours in induce feeding behavior. They were then transferred to the oxidative stress media and assayed for survival regularly until complete lethality, similar to starvation survival.

### Triglyceride Quantification

For each replicate, 5 animals were washed in cold PBS before being homogenized in 500μL PBS + 0.05% Tween20 (PBST) in a microcentrifuge tube using a plastic pestle. 20μL of homogenate was set aside for Bradford protein assay. Remaining homogenate was heated for 10 minutes at 70°C to kill enzymatic activity. 20μL of heated homogenate each was set aside into two microcentrifuge tubes: a reagent tube and a background tube. To the reagent tube, 20μL of Triglyceride Reagent (Sigma) was added to cleave hydrocarbon tails from glycerol. To the background tube, 20μL of PBST was added. All tubes were incubated for 60 minutes at 37°C to promote enzymatic activity. Afterwards tubes were centrifuged for 3 minutes at 10,000 rpm. A standard curve ranging from 0.0625-1.0 mg/mL glycerol standard (Sigma) was diluted from stock into PBST. (The standard curve included points at 0.0625, 0.125, 0.25, 0.5, and 1.0 mg/mL.) 30uL of each standard (including blank) and sample was transferred to a 96-well plate. For colorimetric assay, 100uL Free Glycerol Reagent (Sigma) was added to each well and the plate was incubated at 37°C for 5 minutes to promote enzymatic activity. Absorbance was measured at 540nm on a BMG Labtech POLARstar® Omega plate reader.

For Bradford protein assay, a standard curve ranging from 0.094-1.5 mg/mL bovine serum albumin (BSA) was diluted from stock into ddH2O. (The standard curve included points at 0.09375, 0.1875, 0.375, 0.75, and 1.5 mg/mL.) 10μL of each standard (including blank) was transferred to a 96-well plate. For each homogenized sample, 10μL was transferred to one well and 5μL was diluted in 5μL ddH2O in a second well to test linearity and ensure fit on standard curve. 300μL “Coomassie Plus – The Better Bradford Assay” reagent (Thermo) added to each well and the plate was allowed to sit at room temperature for 10 minutes. Absorbance was measured at 595nm on a BMG Labtech POLARstar® Omega plate reader.

TAG concentration for each replicate was calculated by calculating the concentration of each sample well using the standard curve, and then subtracting the background tube concentration from the reagent tube concentration. Protein concentration for each replicate was calculated by calculating the concentration of each sample well using the standard curve. Average animal TAG levels were calculated by dividing TAG concentration by protein concentration for each replicate and then dividing by 5 (for each animal per replicate).

## Results

Drosophila have a single MLR COMPASS-like complex containing essential subunits Trr and Cmi. Trr harbors the methyltransferase enzymatic activity, and loss of Trr reduces complex stability and prevents formation ^4,51^. Cmi (also known as Lpt) includes histone binding domains. While the complex remains stable and localizes to target enhancer in the absence of Cmi, essential functions such as H3K4me1 deposition and p300 recruitment are lost ^3,49^. To investigate MLR complex function in the Drosophila fat body, we chose to modify complex activity via alteration of Cmi expression. Not only has alteration of Cmi levels been previously demonstrated to directly alter MLR complex activity, but Cmi modulation also often leads to less severe phenotypic changes than Trr modulation, providing an ideal model for phenotypic comparison ^3,4,12^. To accomplish this a Cmi-specific hairpin or full-length Cmi cDNA was expressed specifically in adipocytes by the Lsp-Gal4 driver. Neither fat body-specific Cmi knockdown (Cmi KD) nor Cmi overexpression (Cmi OE) resulted in any observable defects in fat body development or function in normal growth conditions. Fat body size and shape was not noticeably affected by Cmi level, and adipocyte morphology was similarly unaltered (Fig. 1A). Triglyceride storage levels at the end of larval development was similar across all genotypes, suggesting that the larval fat body’s main function as an energy reservoir is not sensitive to Cmi (Fig. 1B). Both Cmi KD and Cmi OE animals shared similar developmental timescales with controls, both larval (Fig. 1C), and pupal (Fig. 1D). The only significant effect resulting from fat body Cmi level in unstressed animals occurred in the context of longevity. Newly-eclosed adults were separated by sex and survival tracked until natural death. Cmi OE males died slightly yet statistically significantly sooner (Fig. 1E), while Cmi KD females survived longer (Fig. 1F). Despite inconsistencies between sexes, these data suggest that Cmi in the fat body has a marginally deleterious effect on longevity. Altogether, these results suggest that MLR complex activity is not required to regulate the post-embryonic development and function of fat body adipocytes during normal growth and development. As the Lsp2-Gal4 driver is expressed only after adipocyte identity is determined during differentiation, it remains a possibility that the complex is required for adipogenesis prior to this point.

**Figure 1.**
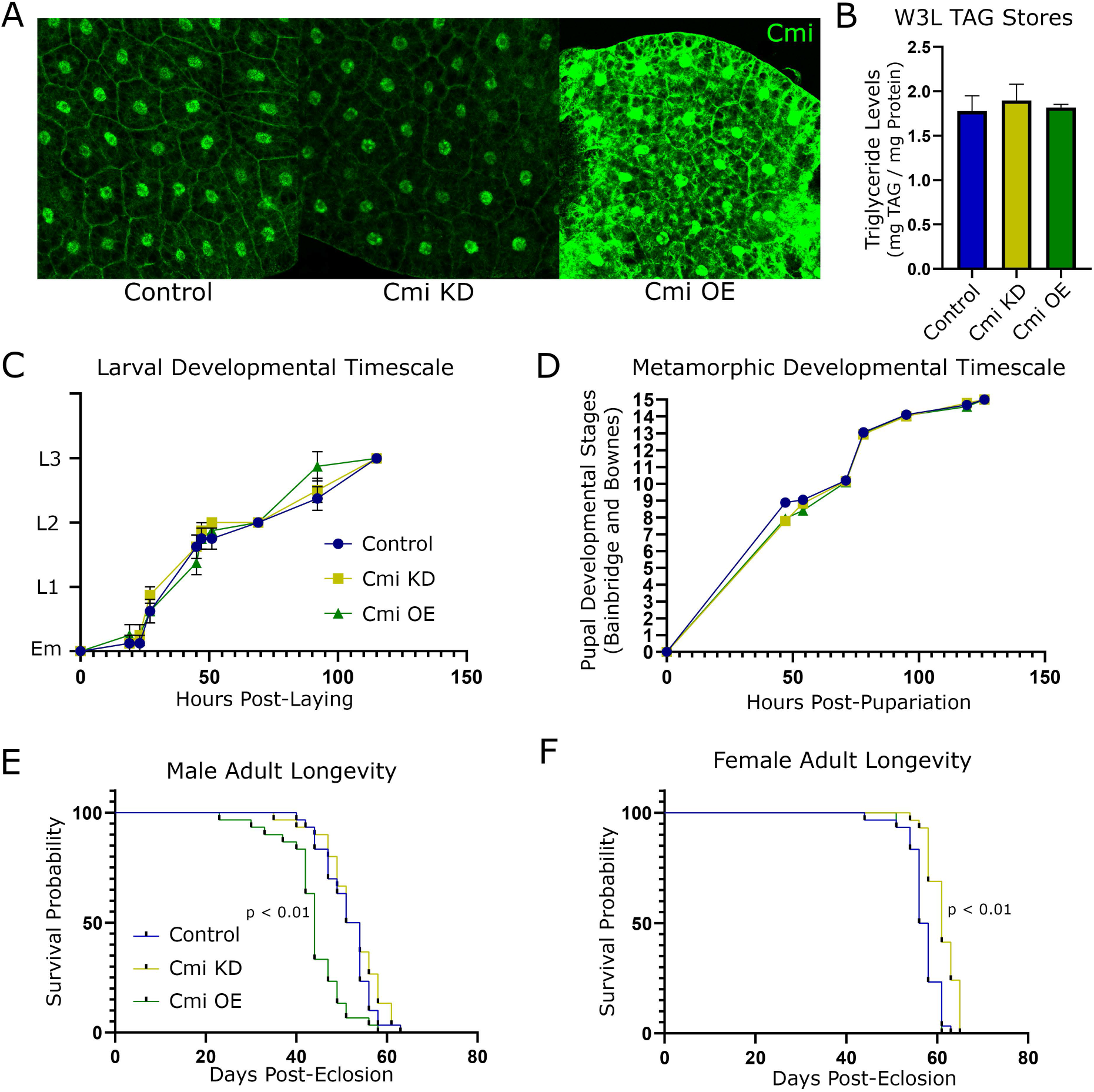
Fat Body Development and Triglyceride Storage is Not Sensitive to Cmi Level. (A) Fat body tissue was dissected from control, Cmi KD, and Cmi OE wandering third instar larvae (W3L) and stained for Cmi. While Cmi level is clearly different among the three genotypes, there is no obvious different in adipocyte size or morphology. (B) W3L were assayed for total stored triglyceride (TAG) content. No significant difference in TAG levels between each genotype was observed. (N = 4, for each genotype; statistical significance measured by Student’s T Test, NS = not significant) (C) Embryos were collected 0-2 hours post-laying and development was visually tracked. Each genotype progressed through hatching and larval development at similar rates. (N = 8 for each genotype; Em=Embryo, L1=first instar larva, L2=second instar larva, L3=third instar larva) (D) W3L were collected just before pupariation and metamorphic development was visually tracked according to Bainbridge and Bownes numeric pupal staging ^50^. Each genotype progressed through hatching and larval development at similar rates. (N = 19 for each genotype) (F-G)

While apparently dispensable during TAG synthesis and storage, the MLR complex may play an important role during TAG depletion. Throughout the process of metamorphosis, the pupa has no access to environmental nutrients and must therefore rely on energy accumulated during the larval period mainly stored as triglycerides in the fat body ^25^. Post-eclosion, remaining triglyceride stores remain in the remodeled adult fat body and are catabolized during the initial non-feeding period of the adult. To investigate if MLR complex activity is required for TAG depletion from the fat body, survival of Cmi KD and Cmi OE pupae undergoing metamorphosis was tracked. At 25°C all genotypes demonstrated similar eclosion rates and displayed no physical differences, suggesting that Cmi level in the fat body has no effect on the animal’s ability to successfully form an adult (Fig. 2A). Drosophila metabolic and developmental rates are directly related to environmental temperature; at the stressful temperate of 29°C, pupa undergo metamorphosis at an accelerated rate yet burn through a greater total amount of triglyceride stores ^25^. At 29°C no Cmi OE animals survived metamorphosis, failing to initiate eclosion and dying as a fully formed pharate inside the pupal case (termed “late pupa lethal”), demonstrating a severe sensitivity to Cmi level at higher temperatures (Fig. 2B). It is possible that Cmi OE pupae are temperature sensitive at a specific metamorphic timepoint, eventually leading to the striking lethality phenotype. If so, identification of that timepoint would provide a clue to the underlying mechanisms. To investigate this, pupae were exposed to two temperature-switch conditions: undergoing the intial half of metamorphosis at 25°C before being transferred to 29°C for latter half, or vice versa (Fig. 2A). Intriguingly, neither condition demonstrated greater lethality of Cmi OE pupae. Instead, while the majority of each cohort displayed late pupal lethality, approximately one quarter began the eclosion process yet died before completion (termed “eclosion lethal”) (Fig. 2C). A small number of Cmi OE pharates were identified alive during the process of eclosion and manually removed from the pupal case. These adults appeared sluggish and were transferred to food, surviving for approximately two hours before dying of unknown causes. These results demonstrate that Cmi OE animals are not dying due to inability to initiate or complete eclosion, but instead are general susceptible to higher temperature during the entire process of metamorphosis. After the initial stages of pupariation, eclosion is the most energetically taxing process of metamorphosis ^25^. The metamorphic lethality data support a hypothesis that Cmi OE pupae burn through fat stores more rapidly, essentially exhausting energy stores when exposed to metabolically stressful high temperatures. If so, then even the surviving Cmi OE adults at 25°C would demonstrate lower remaining triglyceride levels than control. Indeed, while each genotype commences metamorphosis with similar stored fat levels (Fig. 1B), Cmi OE animals exit the process with significantly less than control (Fig. 2D), suggesting that excess levels of Cmi in the fat body increase the triglyceride depletion rate during metamorphic transformation.

**Figure 2.**
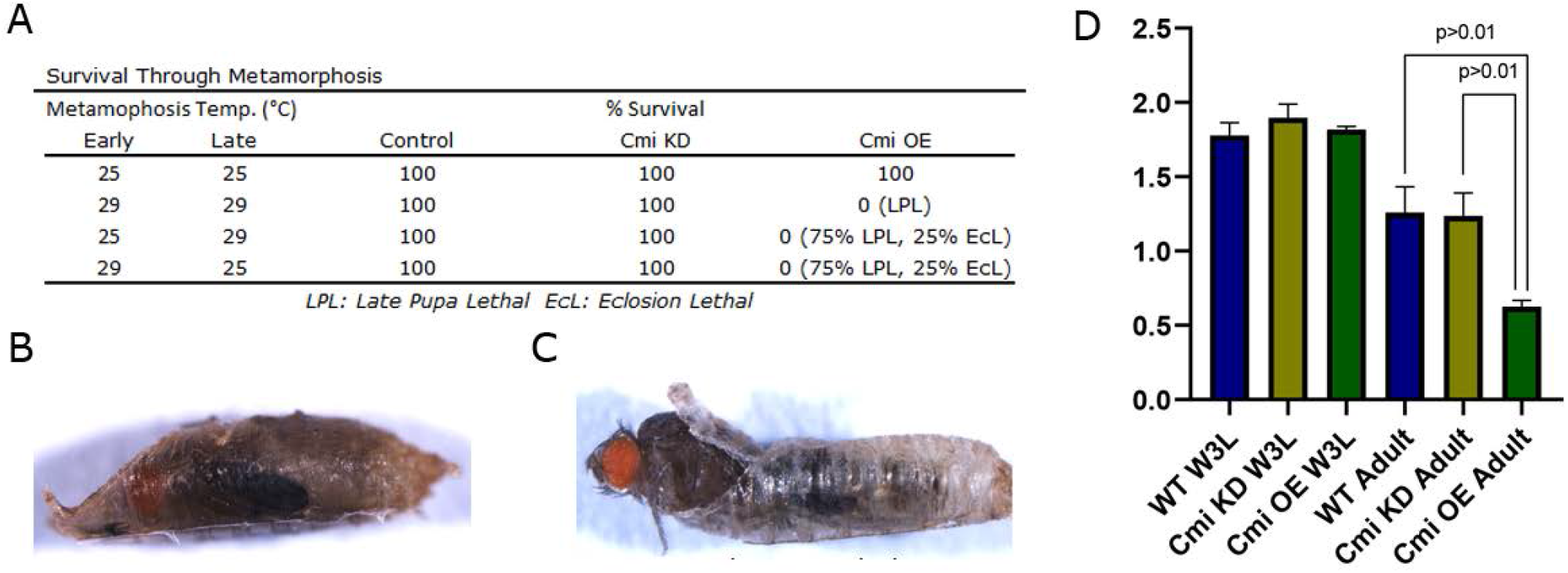
Cmi Overexpression in the Fat Body Causes Increased Triglyceride Depletion during Metamorphosis. (A) Pupae underwent metamorphosis at 25°, 29°, or transferred between the two temperatures. Control and Cmi KD pupae successfully completed metamorphosis and eclosed as healthy adults at all temperatures. At 29°, all Cmi OE pupae died before eclosion and were scored as Pharate Lethal (PL). At either temperature switch condition, approximately 75% of Cmi OE pupae were scored as PL, while the remaining 25% were scored as Eclosion Lethal (EcL). (B) An example of the LPL phenotype, consisting of a fully formed adult that does not begin the eclosion process. (C) An example of the EcL phenotype, consisting of a fully formed adult dying before completion of the eclosion process. (D) Adults having just completed metamorphosis at 25° were assayed for remaining triglyceride (TAG) content. While Cmi KD had no effect on TAG stores compared to control (Fig. 1B), Cmi OE animals demonstrated significantly lower levels. (N = 4, for each genotype; statistical significance measured by Student’s T Test, NS = not significant)

We next sought to determine if Cmi OE in the fat body results in increased triglyceride depletion at other developmental stages or if the effect is specific to metamorphosis. When exposed to nutrient deprivation, adult Drosophila also mobilize adipocyte triglycerides for energy. Newly-eclosed adults were transferred to starvation media (access to moisture but no nutrients) and survival was tracked. As expected, Cmi OE animals, which begin nutrient deprivation with lower fat stores, succumb to starvation significantly sooner than control (Fig. 3A). Surprisingly, despite the fact that Cmi KD adults, which begin starvation with similar triglyceride levels as control, survive significantly longer. Measurement of triglyceride levels during starvation revealed that Cmi KD animals retain significantly higher levels of triglycerides than control flies at both two and three days post-starvation (Fig. 3B). These results suggest that reduced MLR complex activity in the adult fat body suppresses triglyceride depletion rate during starvation.

**Figure 3.**
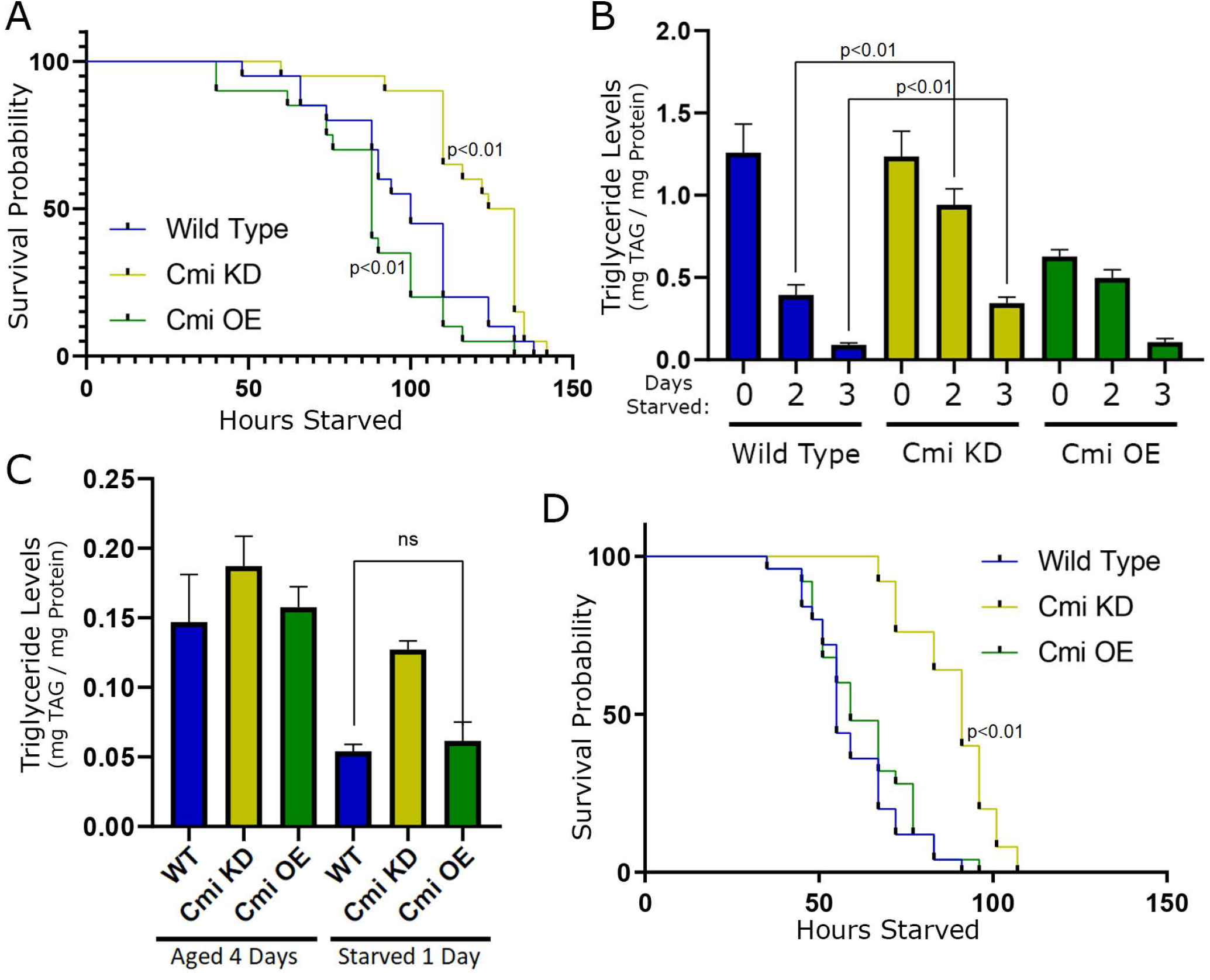
Cmi Knockdown in the Fat Body Causes Decreases Triglyceride Depletion during Adult Starvation. (A) Survival of newly-eclosed adults was tracked during starvation. Cmi KD animals survived significantly longer while Cmi OE animals died significantly sooner. (N = 25, for each genotype; statistical significance measured by Log-rank Test) (B) TAG levels were assayed from representative cohorts exposed to starvation. Cmi KD animals depleted TAGs at a significantly lower rate than control. The depletion rate of Cmi OE animals cannot be compared to control, as the two populations begin starvation with different stored levels. (N = 4 for each genotype; statistical significance measured by Student’s T Test) (C) Newly-eclosed adults were aged 4 days to reach a homeostatic adult TAG level before being starved for one day, and TAG levels were assayed in representative cohorts from each timepoint. After 1 day of starvation, depleted TAG stores of Cmi OE animals were similar to control. (N = 4, for each genotype; statistical significance measured by Student’s T Test, NS = not significant) (D) Adults aged 4 days were starved to lethality and survival tracked. Cmi KD animals survived significantly longer than control, yet Cmi OE animals died at a similar rate to control. (N = 25, for each genotype; statistical significance measured by Log-rank Test, NS = not significant)

The rate of triglyceride depletion during starvation is regulated by multiple factors, including the current level of triglyceride reserves: the less triglycerides available, the lower the rate of depletion ^26^. As Cmi OE adults begin starvation with less fat stores than control, the TAG depletion rate of these animals will necessarily be lower; therefore, any effect of Cmi OE on depletion rate cannot be accurately interpreted from these results. To properly analyze this, all genotypes must begin the process with similar triglyceride stores. Adults of each genotype were collected after eclosion and aged four days until a homeostatic adult triglyceride levels were achieved, which proved to be statistically similar among each genotype (Fig. 3C). After these animals were subjected to one day of starvation Cmi KD animals again demonstrated higher remaining triglyceride levels, yet Cmi OE stores remained similar to control, suggesting that during adult starvation excess Cmi levels do not affect the rate of triglyceride depletion. When starvation lethality was tracked in these aged animals, the sensitivity of Cmi OE animals was abrogated (Fig. 3D). These results demonstrate that while Cmi KD reduces the depletion rate of triglyceride during adult starvation, Cmi OE has no effect.

As the sensitivities to Cmi level on fat body function are specific to developmental stage, it is possible that the MLR complex is required to regulate adult triglyceride storage despite that fact that it is dispensable in the larva. To test this, starved adults were allowed to resynthesize their fat stores on a nutrient-rich medium. After both one and two days of refeeding, Cmi OE and control animals still displayed statistically similar TAG levels, suggesting that excess Cmi in the adult fat body does not affect the ability to synthesize and store triglycerides (Fig. 4). The TAG synthesis rate of Cmi KD adults cannot accurately be compared to control, as synthesis rate (similar to depletion rate) is partially regulated by current TAG level and the two genotypes began refeeding at significantly different levels.

**Figure 4.**
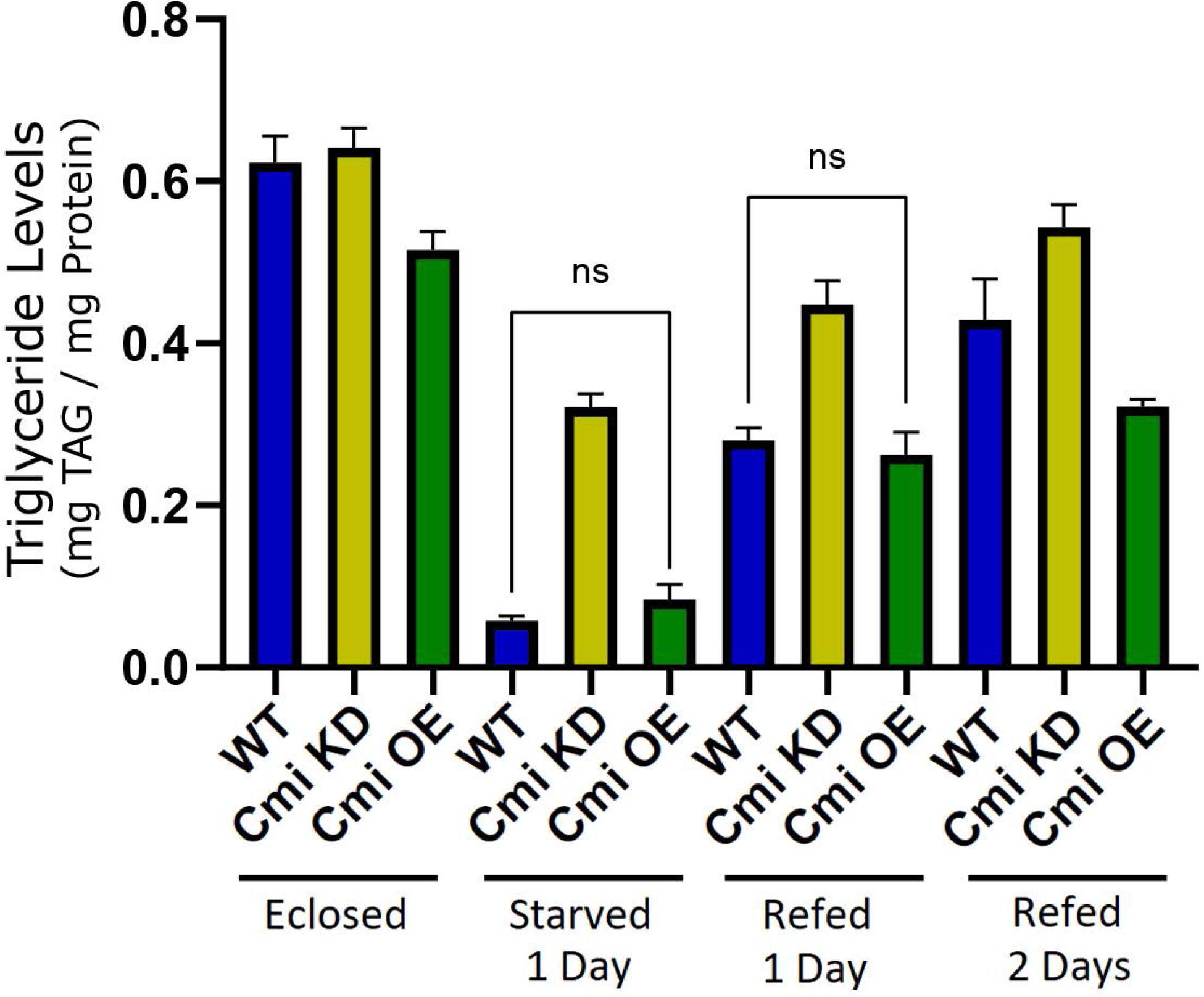
Cmi Overexpression in the Adult Fat Body Has No Effect on Triglyceride Synthesis. Four-day old adults were starved for one day and then refed for two days. TAG levels were assayed in representative cohorts from each timepoint. Cmi OE animals restored TAG levels at a similar rate to control. The TAG replenishment rate of Cmi KD animals cannot be compared to control, as the two populations begin refeeding with different stored levels. (N = 4, for each genotype; statistical significance measured by Student’s T Test, NS = not significant)

These results demonstrate that while the MLR complex appears unnecessary for triglyceride synthesis and storage in adipocytes, it is required to properly regulate the rate of triglyceride depletion in the pupa as well as the adult. The data suggest that in the fat body MLR complex activity serves to promote or enhance the animal’s ability to catabolize stored triglycerides during nutrient stress. However, the fact that the sensitivities to Cmi KD or Cmi OE are developmental stage-specific effects suggests different mechanisms may be at play during different developmental stages. No matter the exact mechanism(s), if the MLR complex is required to regulate triglyceride depletion, it would be through transcriptional regulation. The complex may promote expression of triglyceride mobilization machinery, such as Bmm or Lip3^20^. Metabolic function is also indirectly regulated by hormone and stress response signaling. Ecdysone hormone levels increase during non-feeding periods, and ecdysone receptor (EcR) activation promotes triglyceride mobilization ^52–54^. The MLR complex is recruited by EcR as a coregulator of transcriptional activity ^55^, suggesting that the complex may be required for promoting expression of starvation-associated EcR fat body targets such as Ilp6 or Bmm ^23^.

For an unbiased investigation of changes to gene expression upon knockdown or overexpression of Cmi, we performed RNA-seq analysis on Drosophila fat body tissue. We sought to characterize gene dysregulation occurring during developmental reprogramming, particularly when transitioning to a period of non-feeding. Therefore, we collected fat body tissue from two adjacent developmental stages: the wandering third instar larva “blue gut” (BG), and the “clear gut” (CG). The BG stage occurs just prior to a major ecdysone hormonal spike triggering reprogramming in preparation for the commencement of metamorphosis, while the CG stage occurs immediately after the spike and just prior to pupariation. We anticipated that the results of comparative analyses would demonstrate that modulation of Cmi level in the fat body causes dysregulation of critical metabolic pathways regulating the rate of triglyceride depletion.

Unexpectedly, gene ontology (GO) analysis did not identify lipid metabolic processes as significantly affected upon knockdown or overexpression of Cmi. Instead, Cmi modulation in the fat body results in significant dysregulation of stress response genes, including those activated in response to heat, oxidative, and antimicrobial stresses (Fig. 5A). Manual interrogation of the RNA-seq results revealed multiple patterns of transcriptional dysregulation among stress response genes, including AMPs and other Foxo targets.

**Figure 5.**
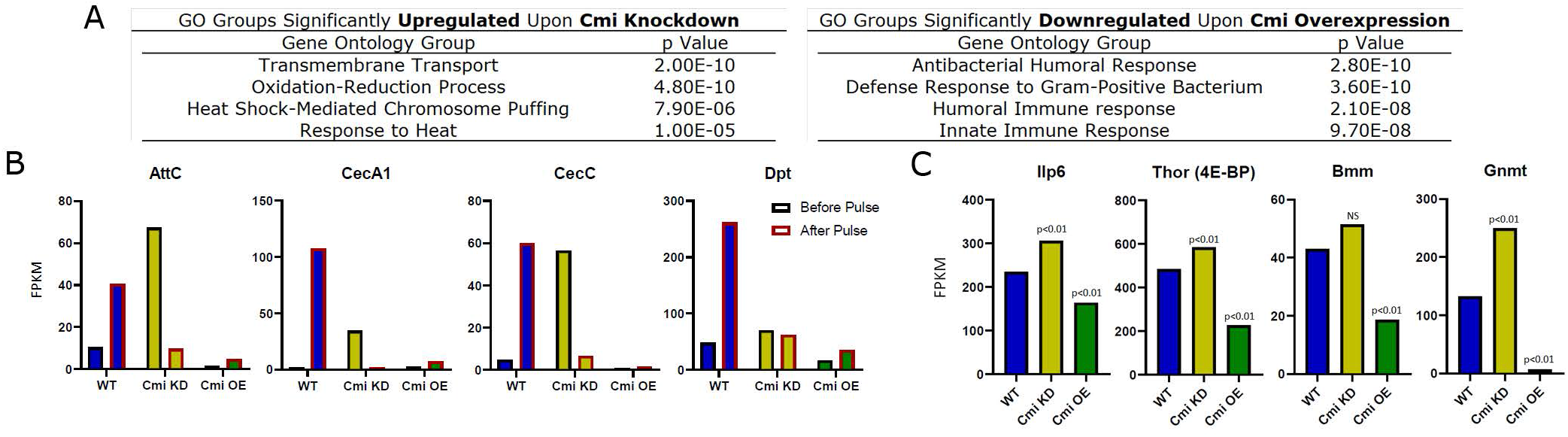
Modulation of Cmi Levels in the Larval Fat Body Significantly Dysregulates Stress Response Transcription. RNA-seq was performed on larval fat bodies to identify differential gene expression profiles associated with Cmi KD and Cmi OE. (A) Gene ontology (GO) analysis demonstrates that stress response expression is significantly dysregulated upon alteration of Cmi levels, as illustrated by the top four most significantly altered GO categories upregulated in blue gut larvae upon Cmi KD (left) and downregulated in blue gut larvae upon Cmi OE (right). (B) In wild type fat body, expression of antimicrobial peptides (AMPs) AttC, CecA1, CecC, and Dpt is upregulated upon transition from BG to CG stage. Upon Cmi KD, AMP expression is increased in the BG and decreased in the CG compared to control. Upon Cmi OE, AMP expression in decreased in both the BG and CG compared to control. (C) Foxo transcriptional targets in the fat body (Thor, Ilp6, Bmm, and Gnmt) are all upregulated upon Cmi KD and downregulated upon Cmi OE in the blue gut stage. (Statistical significance determined by Novogene Genomics Services, NS = not significant)

AMPs are expressed and secreted systemically by the fat body in response to pathogenic signatures. AMPs are normally transcribed at low levels in the BG stage and are significantly upregulated during the reprogramming transition to CG (Fig. 5B). Cmi OE larvae simply fail to properly upregulate in CG. However, upon Cmi KD, the normal relationship is reversed: expression is atypically high in the BG yet is downregulated upon transition to CG. This pattern is strikingly similar to the effects of MLR complex loss at EcR target genes that our lab has recently characterized ^3^. The MLR complex is required not only for readying these targets for later activation, but also for suppressing premature activation. These results suggest that the complex plays a similar role in the fat body at innate immunity genes.

Foxo, the highly-conserved stress-response facto, is not only responsible for AMP expression during metamorphosis, but also drives expression of multiple genes upregulated during non-feeding periods; these include TAG lipase Bmm, system growth regulator Ilp6, translation regulator Thor (4E-BP), and energy homeostasis regulator Gnmt ^40,56^. These fat body regulatory targets of Foxo are all upregulated upon Cmi KD and downregulated upon Cmi OE (Fig. 5C), suggesting that the MLR complex is involved in suppressing Foxo activity in the larval fat body.

Cellular metabolism is often indirectly regulated by stress response transcription, as homeostatic cellular processes are suppressed while addressing the stress state. If the MLR complex is required to properly regulate transcriptional reprogramming in reaction to stress, then the observed triglyceride depletion alterations may be indirect effects of a dysregulated stress response. To investigate this, we used the Gal4 system to either knockdown (Foxo KD) or overexpress (Foxo OE) Foxo in the fat body and subjected the resulting adults to starvation. Given the inverse relationship between Cmi and Foxo activity suggested by the larval RNA-seq data, we anticipated that Foxo KD would mimic the effects of Cmi OE (starvation sensitivity) and that Foxo OE would mimic Cmi KD (starvation resistance). Unexpectedly, the reverse proved true: Foxo KD provided a significant protective effect similar to Cmi KD, while Foxo OE animals succumbed to starvation significantly sooner, similar to Cmi OE (Fig. 6A). Additionally, just as in Cmi OE animals, aging Foxo OE adults before nutrient deprivation abrogated the susceptibility to starvation lethality (Fig. 6B). These results suggest that the MLR complex promotes Foxo activity in the adult fat body.

**Figure 6.**
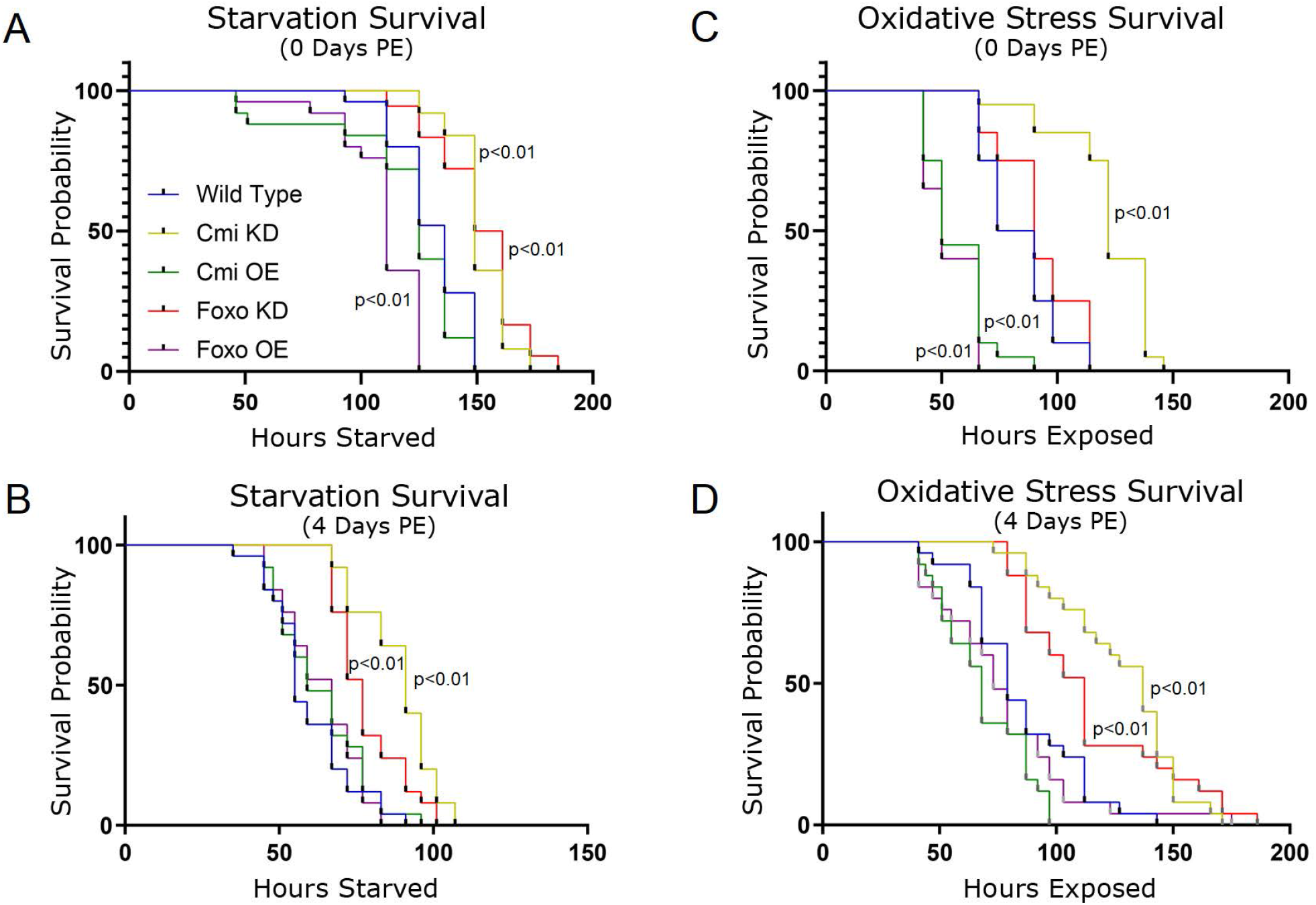
Modulation of Foxo Levels in the Adult Fat Body is Phenotypically Similar to Modulation of Cmi. (A) When newly-eclosed adults are starved, Cmi KD and Foxo KD animals survive significantly longer while Cmi OE and Foxo OE animals die sooner. (B) When four-day old adults are starved, Cmi KD and Foxo KD animals survive significantly longer while Cmi OE and Foxo OE has no effect on survival. (C) When newly-eclosed adults are exposed to oxidative stress via H2O2, Cmi KD and Foxo KD animals survive longer while Cmi OE and Foxo OE animals die significantly sooner. (D) When four-day old adults are exposed to oxidative stress via H2O2, Cmi KD and Foxo KD animals survive significantly longer while Cmi OE and Foxo OE animals die sooner. (N = 25, for each genotype; statistical significance measured by Log-rank Test, NS = not significant)

If the MLR complex is necessary for Foxo transcriptional activity in the adult fat body, then processes requiring Foxo would also require activity of the MLR complex. Foxo is activated in response to oxidative stress, and loss of Foxo activity inhibits the animal’s ability to survive this stress ^39,56^; the role of Foxo specifically in the fat body in response to oxidative stress, however, is less certain ^57^. Newly-eclosed adults either modulating Cmi or Foxo levels in the fat body were exposed to lethal levels of hydrogen peroxide (H_2_O_2_), a reactive oxidant. Both Cmi KD and Foxo KD significantly extended survival of these animals compared to control, Cmi OE and Foxo OE significantly reduced survival (Fig. 6C). These results support the theory that the MLR complex is required to promote Foxo activity in the adult fat body, and further suggest that Foxo activity in this organ is somehow detrimental to the animal’s survival in response to stress. Notably, whereas aging of Cmi OE and Foxo OE animals nullifies susceptibility to starvation (likely by overcoming triglyceride deficiency after metamorphosis), aged Cmi OE and Foxo OE adults retain their susceptibility to oxidative stress (Fig. 6D). This suggests that multiple mechanisms affecting survival are regulated by Foxo and the MLR complex.

## Discussion

In recent years, MLR complexes have become well-characterized as enhancer regulators required for proper transcriptional reprogramming in response to signaling factors and other developmental cues during cellular differentiation and tissue patterning. MLR complexes also play necessary regulatory roles after development, including being required in terminally-differentiated cells for transcriptional responses to homeostatic maintenance cues as coregulators of FXR and p53 targets ^16,58–60^. The results presented here identify further examples of this latter type of role, suggesting that the Drosophila MLR complex is necessary for controlling the depletion rate of fat body TAGs during nutrient stress, a precisely regulated function critical for organismal survival.

Evidence during metamorphosis and adult starvation suggest that in the fat body the MLR complex plays a role in suppressing TAG depletion during nutrient stress. However, the effects are stage-specific: Cmi KD inhibits the rate of TAG depletion during adult starvation yet has no effect during metamorphosis; Cmi OE significantly enhances TAG depletion during metamorphosis but has no effect during starvation. As the complex is required for the proper regulation of gene expression before and during reprogramming events, these stage-specific effects may be due to the dysregulation of different transcriptional targets and/or the involvement of different regulatory machinery (e.g., transcription factors) during each developmental stage.

Transcriptome analysis determined that the expression of stress response genes is significantly sensitive to Cmi levels in the fat body, suggesting that a major role of the MLR complex in this tissue is regulating transcriptional reprogramming in response to stress signals. This hypothesis was supported by oxidative stress survival assays, which demonstrated an inverse relationship between Cmi level and survival. These results suggest that reduction of MLR complex activity in the adult fat body protects against oxidative stress, just as it does during starvation.

The cellular metabolic state is indirectly regulated by stress response pathways; this feeding/stress axis is particularly critical in the fat body, a regulator of both systemic growth and stress response. Our larval fat body transcriptomic data suggests that the MLR complex functions through suppression of the targets of Foxo, a highly conserved transcription factor essential for stress response reprogramming. This is challenged by the fact that there is positive correlation between Cmi and Foxo levels in the fat body during both starvation and oxidative stress survival, suggesting a positive mechanistic relationship between the two. While the transcriptomic and survival data appear conflicting, we have previously established that the MLR complex may have opposite regulatory activity on the same transcriptional targets at different developmental timepoints ^3,4^. Therefore, we posit that the MLR complex is responsible for suppressing transcription of Foxo targets in the larval fat body, yet later serves to enhance expression of Foxo targets in the adult fat body.

We have suggested that this form of developmental differential regulation is due to dual regulatory roles of the MLR complex: suppression of premature transcription prior to activating signals, and subsequent enhancement of gene activation in the presence of those signals ^3^. In the fat body we observe a similar pattern in the expression of AMPs, which are normally significantly upregulated during the transition of blue gut to clear gut prior to pupariation. The fact that this relationship is reversed upon loss of complex activity, in a relationship similar to previously characterized ecdysone response genes, suggests that the MLR complex plays the same dual regulatory role at these genes, likely acting as a coregulator with one or more of the Drosophila NF-κB-like effectors: Relish, Dl, and Dif.

Under nutrient stress, stress factors are required to properly fine-tune the rate of TAG depletion via transcriptional regulation lipase Bmm: Foxo activity enhances depletion through promotion of Bmm expression, whereas Relish activity suppresses Bmm transcription ^43^. Based on the evidence presented here, we propose that the MLR complex is required to inhibit the rate of TAG depletion during non-feeding periods via promotion of Foxo activity and/or suppression of Relish activity (Fig. 7). These effects may be caused by MLR complex regulation at multiple levels, including transcription of Foxo/Relish, regulation of upstream modifiers/signaling machinery, and coregulation of downstream targets.

**Figure 7.**
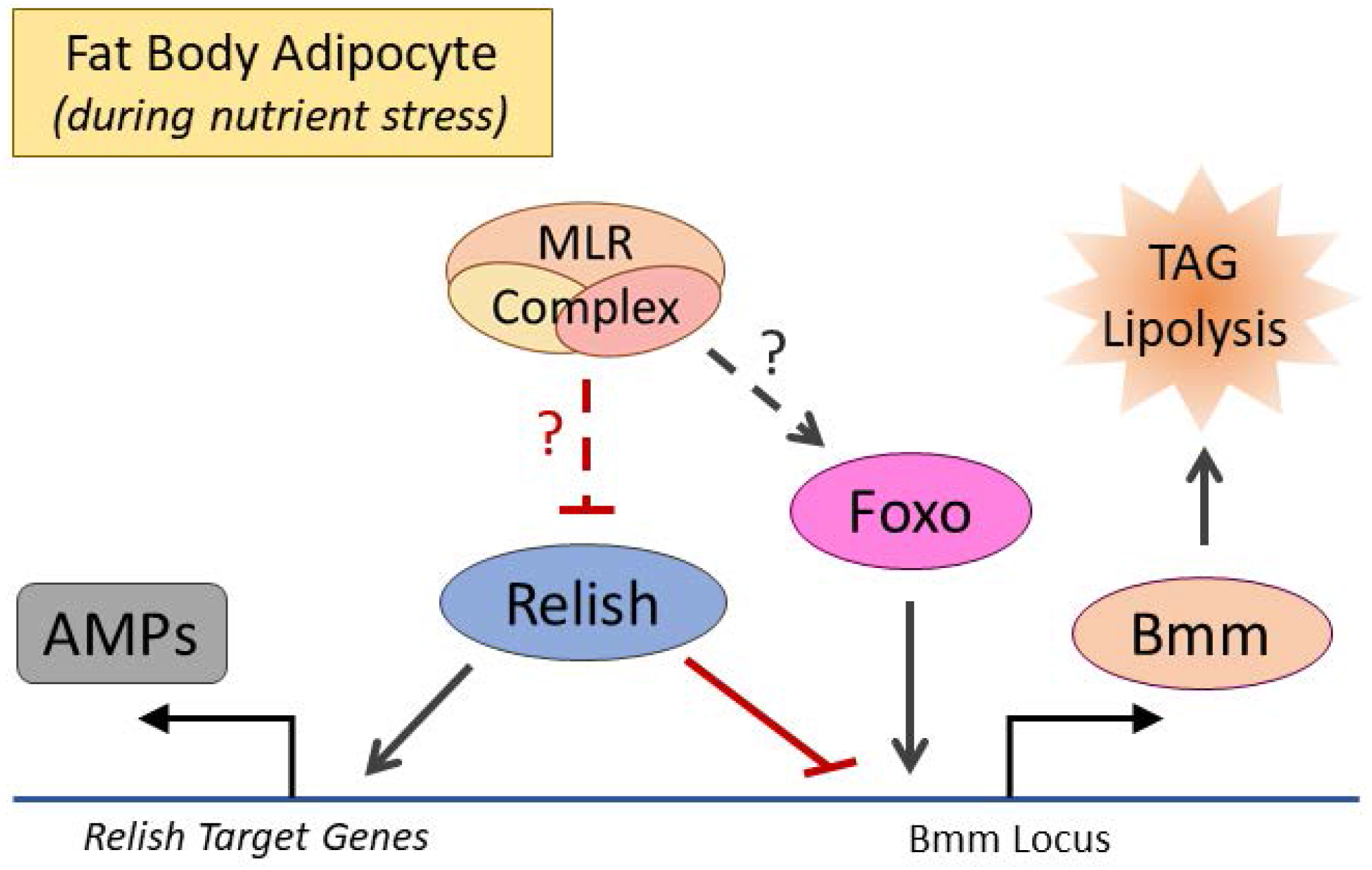
Model. The results presented here suggest that MLR complex activity promotes TAG depletion during nutrient stress and is required to regulate targets of stress response effectors Foxo and Relish. TAG lipolysis is regulated by Bmm transcription, which is promoted by Foxo and inhibited by Relish. We propose that, in times of nutrient stress, the MLR complex is required to positively regulate Foxo activity and/or negatively regulate Relish activity, thereby controlling the rate of TAG depletion. The exact mechanisms of regulation, including whether the MLR complex is required upstream or at the level of Bmm transcription, are unknown.

This report further elucidates the roles of MLR complexes during reprogramming in terminally differentiated cells in response to environmental stimuli, an understudied area. These results also suggest new mechanisms that may be involved in human tumors with high rates of MLR methyltransferase mutation. Dysregulation of stress signaling and inflammatory pathways, particularly those involving Foxo and Relish-orthologous effectors, promote survival and metastasis of tumor cells in addition to increasing resistance to treatment ^61–64^. If the requirement of MLR complex activity for proper stress factor-associated reprogramming is conserved in humans, then the alteration of MLR complex activity has clear oncogenic potential.

